# Voting-based integration algorithm improves causal network learning from interventional and observational data: an application to cell signaling network inference

**DOI:** 10.1101/2020.02.18.955153

**Authors:** Meghamala Sinha, Prasad Tadepalli, Stephen A. Ramsey

## Abstract

In order to increase statistical power for learning a causal network, data are often pooled from multiple observational and interventional experiments. However, if the direct effects of interventions are uncertain, multi-experiment data pooling can result in false causal discoveries. We present a new method, “Learn and Vote,” for inferring causal interactions from multi-experiment datasets. In our method, experiment-specific networks are learned from the data and then combined by weighted averaging to construct a consensus network. Through empirical studies on synthetic and real-world datasets, we found that for most of the larger-sized network datasets that we analyzed, our method is more accurate than state-of-the-art network inference approaches.

## Introduction

Causal modeling is an important analytical paradigm in action planning, predictive applications, research, and medical diagnosis [1,2]. A primary goal of causal modeling is to discover causal interactions of the form *V_i_* → *V_j_*, where *V_i_* and *V_j_* are observable entities and the arrow indicates that the state of *V_i_* influences the state of *V_j_*. Causal models can be fit to passive observational measurements (*“seeing”*) as well as measurements that are made after performing external interventions (*“doing”*).

In many settings, observational measurements [3] are more straightforward to obtain than interventional measurements, and thus observational datasets are frequently used for causal inference. However, given only observational data, it is difficult to distinguish between compatible Markov equivalent models [4,5]. For example, the three causal models *V_i_* → *V_j_* → *V_k_*, *V_i_* ← *V_j_* ← *V_k_*, and *V_i_* ← *V_j_* → *V_k_* are Markov equivalent—each encodes the conditional independence statement *V_i_* ╨ *V_k_* | *V_j_*. This ambiguity can in principle be resolved by incorporating measurements obtained from interventional experiments in which specific entities are targeted with perturbations. With the benefit of interventional measurements, Markov equivalent causal models can have different likelihoods, enabling selection of a maximum-likelihood model. These considerations have motivated the development of network learning approaches that are specifically designed to leverage mixed observational and interventional datasets [6].

Learning a causal network from a mixed observational-interventional dataset poses methodological challenges, particularly in integrating datasets from different experiments and accounting for interventions whose effects are uncertain [7]. Due to batch effects, data collected from two different experiments might not be identically distributed and thus the two experiments may be incoherent from the standpoint of causal network model. As a result, directly combining data from different experiments can lead to errors in network learning. Interventions, too pose a challenge due to the fact that in real-world settings many interventions are (i) imperfect, meaning interventions are unreliable and have soft-targets (A “soft” target intervention, or “mechanism change,” is an intervention that changes a target node’s distribution’s parameters, but does not render that it’s independent of its parent nodes [7]), and (ii) uncertain, meaning that the“off-target” nodes are unknown. Classical causal learning algorithms are based on the assumption that interventions are *perfect* [1]; applying such algorithms to a dataset derived from imperfect interventions would likely yield spurious interactions. Eberhardt [8]classifies such errors into two types: a) *independence to dependence* errors, where two variables *V_i_* and *V_j_* that are independent are detected as dependent when data from the observational and interventional experiments are pooled (i.e., false positive detection of a causal interaction) and b) *dependence to independence* errors, where two variables *V_i_* and *V_j_*, that are dependent in an observational study are independent when the data from the observational study are pooled with data from an interventional study (i.e., a false negative for the interaction). Consensus has yet to emerge on the question of how—given two or more datasets generated from different interventions—the datasets should be combined to minimize such errors in the learned network model.

In this paper, we have demonstrated in details the performance of our proposed method, “Learn and Vote”[9], for inferring causal networks from multi-experiment datasets. “Learn and Vote” can be used to analyze datasets from mixed observational and interventional studies and it is compatible with uncertain interventions. As it is fundamentally a data integration method, “Learn and Vote” is compatible with a variety of underlying network inference algorithms; our reference implementation combines “Learn and Vote” data integration with the Tabu search algorithm [10] and the Bayesian Dirichlet uniform (BDeu) [6,11,12] network score, as described below. To characterize the performance of “Learn and Vote”, we empirically analyzed the network learning accuracies of “Learn and Vote” and six previously published causal network learning methods (including methods that are designed for learning from heterogeneous datasets) applied to six different network datasets. Of the six network datasets, the largest real-world dataset is a cell biology-based, mixed dataset (the Sachs et al. dataset [13]) with a known ground-truth network structure. On larger networks, we report superior (or in worst case, comparable) performance of “Learn and Vote” to the six previously published network inference methods.

## Motivation and Background

### Spurious dependencies and independencies

In this section, we introduce notation and describe how perturbations affecting two or more variables in a causal model can lead to spurious dependencies or independencies. Mathematically, a causal model *M_c_* is described by a directed acyclic graph (DAG) containing a pair (*V, E*), where *V* is a set observable nodes (corresponding to random variables), *E* is a set of directed edges between pairs of nodes, Pa(*V_i_*) represents the set of parent nodes of variable *V_i_*, and *P*(*V*) represents the joint probability distribution. In the context of network learning from interventional data, it is helpful to picture an intervention (say, *I*_1_) as a separate type of node (denoted by a dashed circle in Fig. 1) that can be connected to its targets (say, *V_i_* and *V_j_*) by causal edges of a separate type (dashed arrow in Fig. 1). Applying classical network inference algorithms to measurements pooled from multiple interventional experiments can lead to two different types of learning errors, as we explain below.

**Fig 1.**
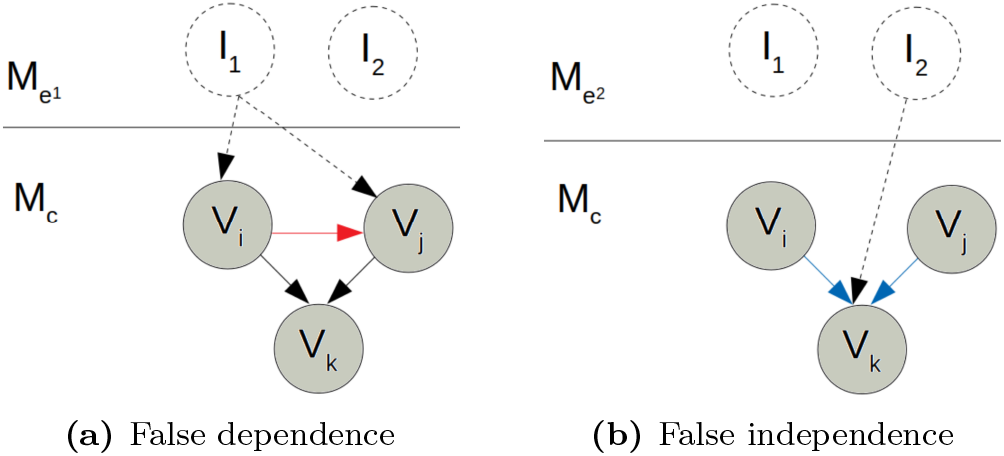
Cross-experiment data pooling leads to network inference errors. Illustration of a simple hypothetical causal model *M_c_* with three observable entities (*V_i_*, *V_j_*, and *V_k_*. Two different interventional experiments are depicted: experiment *M*_*e*^1^_ involves intervention *I*_1_, and experiment *M*_*e*^2^_ involves intervention *I*_2_. Pooling measurements from the two experiments can cause two types of network inference errors: false positive edge (shown in (a) as a red arrow between *V_i_* and *V_j_*), and false negative edges (shown in (b) as blue arrows between *V_i_* and *V_k_* and between *V_j_* and *V_k_*).

1. **False causal dependence:** In the experiment depicted in Fig. 1a, *V_i_* and *V_j_*, which are not causally related in 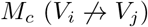, are affected by intervention *I*_1_. Due to the intervention’s confounding effect, we have 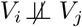 in the combined model *M*_*T*_1__ = *M_c_* + *M*_*e*_1__ (we denote the joint distribution in the combined model by *P*_1_(V ⊂ *M*_*T*_1__). Thus, pooling data from such different distributions may lead to spurious correlations between independent variables.
2. **False causal independence:** In the experiment depicted in Fig. 1b, the intervention *I*_2_ on *V_k_* removes all the incident arrows for *V_k_* and cuts off the causal influences of *V_i_* and *V_j_* on *V_k_*, causing *V_i_* ╨ Pa(*V_i_*). Pooling data from such models can cause the causal dependencies *V_i_* → *V_k_* and *V_j_* → *V_k_* in *M_c_* to be missed (i.e., a “false negative” in the inferred network).

### Review of prior literature

Classical causal learning methods fall into two classes: *constraint-based* methods (e.g., PC [2], FCI [14]), in which the entire dataset is analyzed using conditional independence tests; and *score based* methods (e.g., GES, GIES [15]), in which a score is computed from the dataset for each candidate network model. Both classes of methods were designed to analyze a single observational dataset, with the attendant limitations (in the context of multi-experiment datasets) that we described above. Several multi-dataset network inference approaches have been proposed that circumvent the above-described problems associated with cross-experiment measurement pooling. Cooper and Yoo [6] proposed a score-based algorithm that combines data from multiple experiments, each having perfect interventions with known targets. The approach was later refined by Eaton and Murphy [7] for uncertain and soft interventions [16]. The method of Claassen and Heskes [17] is based on imposing the causal invariance property across environment changes. Sachs et al. [13] analyzed a molecular biology dataset (which has since become a benchmark dataset for molecular network inference, a primary application focus of our work) using a variant of the Cooper-Yoo method. Chen et al. [18] proposed a subgraph-constrained approach, called Trigger, to learn a yeast gene regulatory network model from transcriptome and genotype data. In the Joint Causal Inference (JCI) [19] method, additional experimental context variables are introduced before data pooling. Notably, the aforementioned methods assume some prior knowledge about the network model. In contrast, our “Learn and Vote” method (see Methods and Datasets) requires no prior knowledge about the network model.

#### Network Combination Methods

Another class of multi-dataset network inference approaches, which we call “network combination” methods, involve learning causal interaction statistics from each experiment followed by integration of the statistics to obtain a single consensus network. For example, in the ION [20] method, locally learned causal networks having overlapping variables are integrated. The constraint-based COmbINE [21] method is based on the estimation of variable-variable dependencies and independencies across separate experiments. The MCI [22] algorithm is a constraint-based method that exploits the ‘local’ aspect of causal V-structures [23]. However, none of these methods produce experiment-specific weighted graphs, instead enumerating experiment-specific partial ancestral graphs that are consistent with the data. In real-world datasets, due to a variety of factors (finite sampling, experiment-specific biases and confounding effects, measurement error, missing data, and uncertain/imperfect interventions), the confidence with which a given causal interaction *V_i_* → *V_j_* can be predicted within a given experiment will in many cases vary significantly from experiment to experiment (and in the case of incomplete measurements, may not be quantifiable at all in a given experiment). Thus, a network combination method compatible with experiment-specific edge weights would seem to offer a distinct advantage in the context of multi-experiment network inference. Furthermore, all of these methods assume that a single underlying causal model accounts for all observed causal dependencies. In real-world settings where experimental conditions change across experiments, this assumption seems unlikely to hold, motivating the need for network inference methods that can (1) score candidate interactions within individual experiment-specific datasets and (2) combine weighted edges from experiment-specific datasets into a consensus network.

### Biological Signaling Networks

A cell signaling network is a type of causal network in which the state of a protein or other biomolecule influences the state of another protein or biomolecule downstream of it (denoted by a directed arc). Such networks are amenable to interventional experiments using molecular agents that target (i.e., activate or inhibit) specific molecules. Sachs et al. [13] used a Bayesian network approach to infer causal interactions among eleven signaling molecules in human CD4+ T-cells. In a series of nine experiments—two observational and seven with specific molecular interventions—they measured the activation levels of eleven phosphorylated proteins and phospholipids by flow cytometry (Figure 2). They found 17 true positive interactions, with 15 that were well-established in the biology literature and two that were supported by at least one study; their inferred network missed three arcs (false negatives) and it had no false positive arcs.

**Fig 2.**
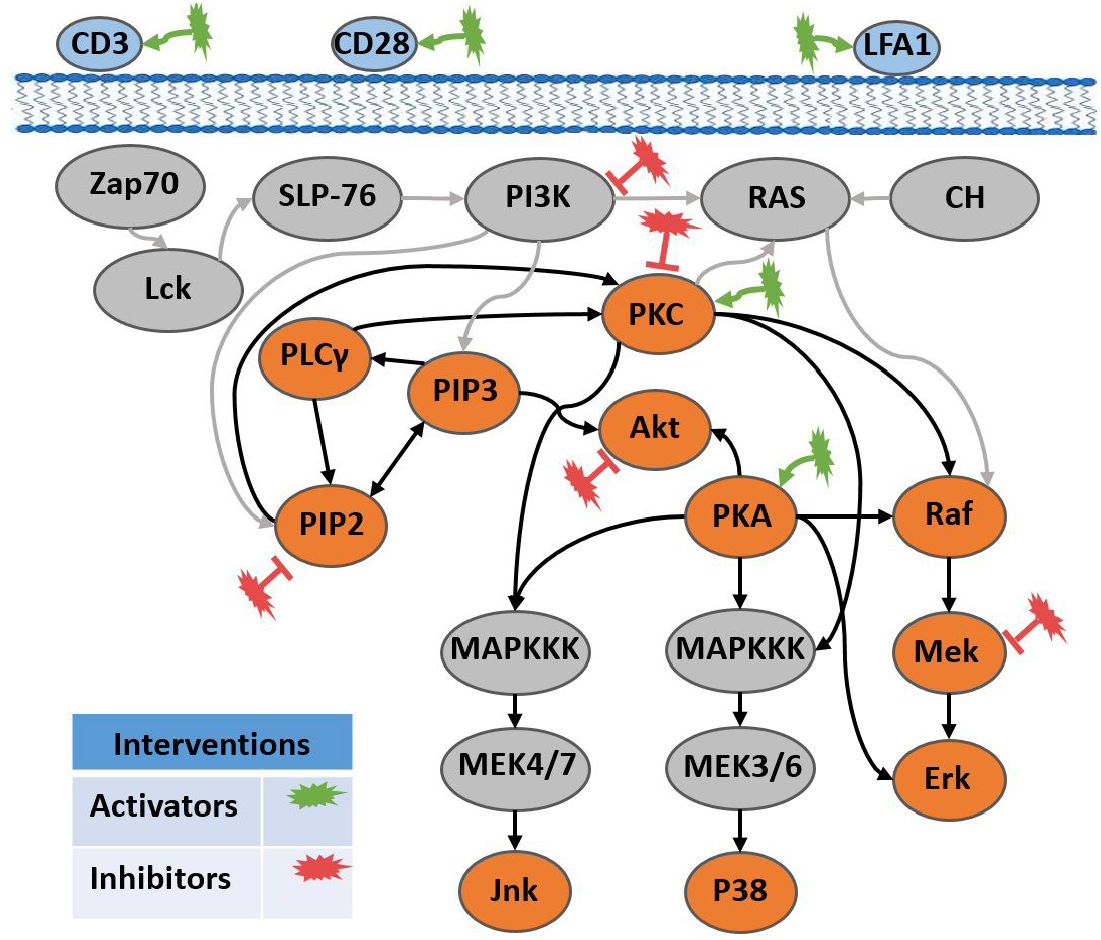
Biological network for the Sachs et al. study, showing interactions (arcs) and interventions (starred ellipses). The pathways represented by bold black lines are the Ground Truth known causal interactions, established through literature study.

### Uncertain interventions

Like most causal network learning approaches, the method used in the Sachs et al. study and in our re-analysis assumes perfect interventions, i.e., that each of the interventional agents targets exactly one of the signaling molecules. Such a perfect intervention assumption is likely not consistent with typical interventions in biological systems, due to potential off-target effects of pharmaceutical agents. Moreover, in a biological system, the effects of certain types of interventions (for example, a gene knockout) may not be describable by forcing of a target node’s state to a specific value in the observational network. In the Sachs et al. experiments, although the interventions are assumed to be perfect, they are known to have off-target effects, as shown by Eaton & Murphy (2007) [7]. Eaton & Murphy modeled chemical interventions as context variables in the network (assuming they had some known background knowledge about the underlying network) to learn the intervention’s effects and found them to have multiple children. To summarize, in the context of current learning algorithms, there are three primary issues with pooling experimental data that were acquired with imperfect interventions:

1. Current algorithms might make mistakes since the arcs pointing towards the unknown targets are not removed or handled properly.
2. Although pooling data adds more confidence into learning the true causal arcs, it can also introduce spurious arcs with incorrect direction (see Fig. 4).
3. Each intervention might alter a mechanism or influence the local distribution in an unknown way [24].

## Methods and Datasets

To avoid the problems arising from pooling data from different experiments in causal network learning, we propose the “Learn and Vote” method (shown in Fig. 3 and Algorithm 1). The method’s key idea is to (1) learn a separate weighted causal network from the data generated in each experiment (which may be interventional or observational) by ignoring the directed arcs into the intervened variables and then (2) combine the experiment-specific networks by weighted averaging. The algorithm’s inputs are, for each experiment, the values of the observed variables (*V*) in the experiments (we denote the number of variables by *v* and the number of experiments by *k*) and the identities of the known target nodes (stored as a list *intv)* for any interventions.

**Algorithm 1.**
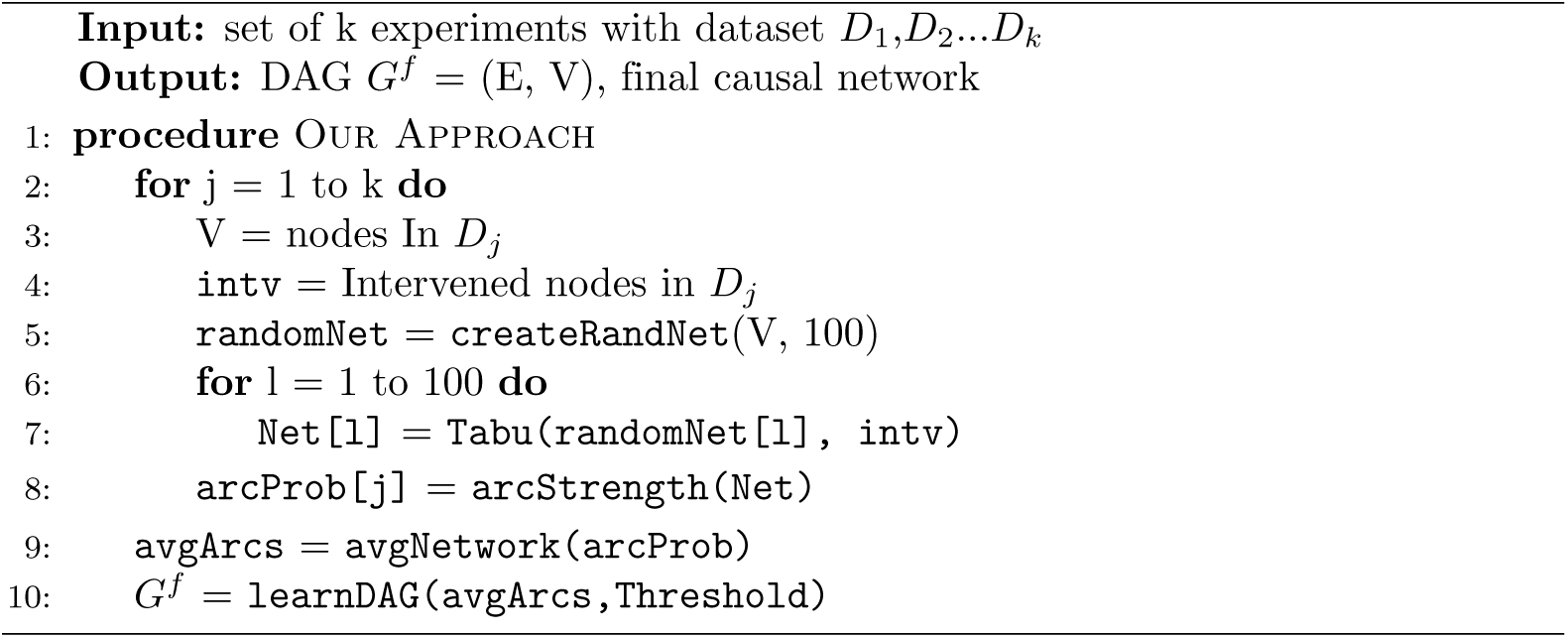
Learn and Vote.

**Fig 3.**
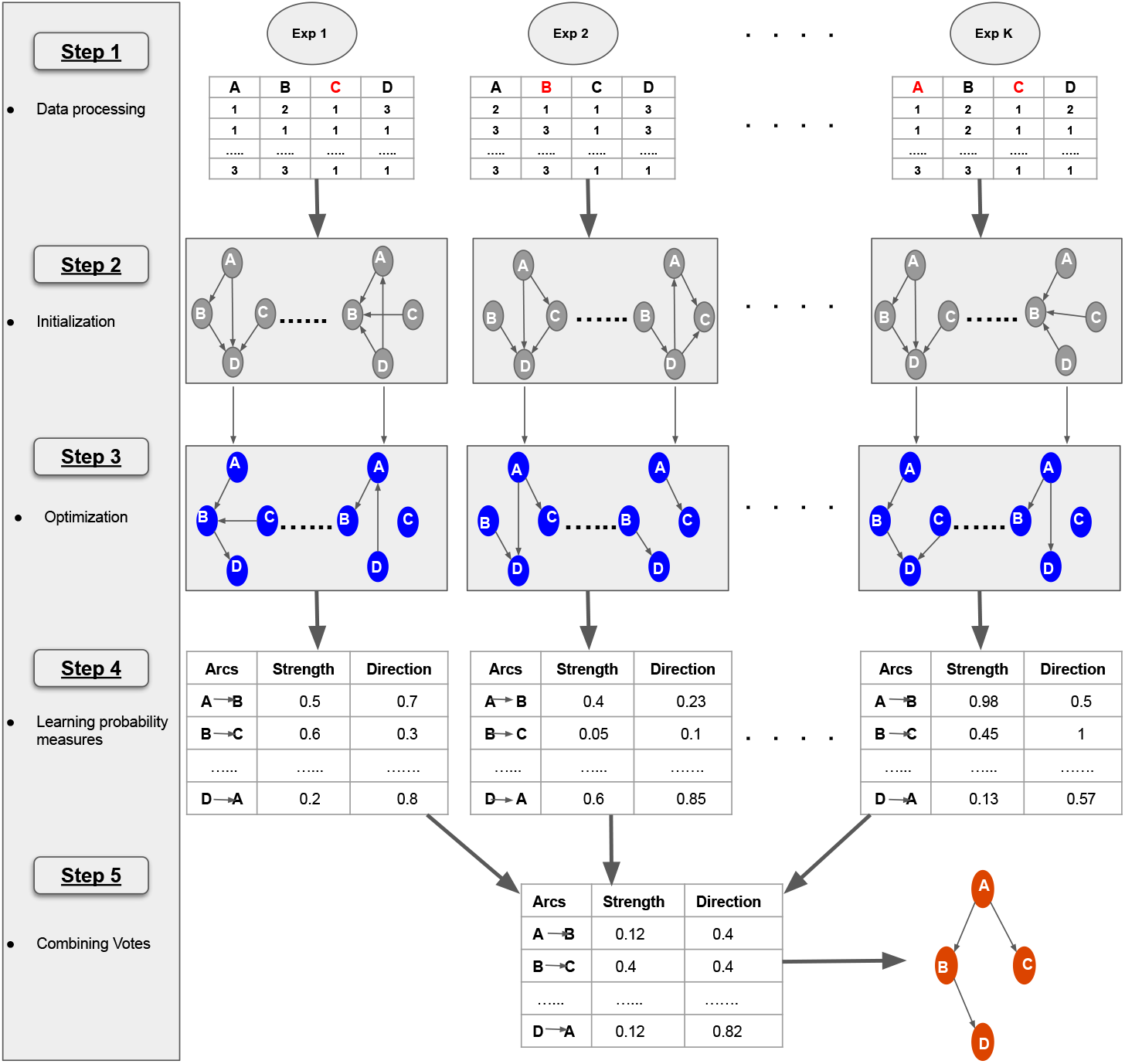
Workflow of “Learn and Vote”: **Step 1** - Collecting data from *k* experiments (combination of observational and interventional studies). For interventional studies, the known targets (marked in Red) are incorporated as external perturbation during the search process. **Step 2** - Creating 100 random DAGs using the observed nodes, as a starting point. **Step 3** - Optimizing each of the 100 DAGs with data using Tabu search. **Step 4** - Calculating probability (in terms of strength and direction) of occurrence for every possible arc from the 100 optimized DAGs and storing them in tables. **Step 5** - Combining votes from all the tables by weighted averaging and constructing the final causal network, with arc strengths above a threshold (in this case 50%)

### Scoring Function

We incorporate the effect of intervention in the score component associated with each node by modifying the standard Bayesian Dirichlet equivalent uniform score (BDeu) [6,11,12]. Given measurements *D_j_* of variables *V* in experiment *j*, let *G^j^* represent a DAG learned from it (with conditional distributions P(*V_i_*|Pa(*V_i_*)*^G^j^^*), where Pa(*V_i_*)*^G^j^^* is the set of parent nodes of *V_i_* in DAG *G^j^*). In a perfect interventional experiment, for the set Int(*m*) of intervened nodes in sample *m*, we fix the values of *V_i_*[*m*] ∈ Int(*m*), meaning that we exclude P(*V_i_*[*m*] | Pa(*V_i_*)[*m*]) from the scoring function for *V_i_* ∈ Int(*m*). All the other unaffected variables are sampled from their original distributions. The distribution of Dj is per experiment and not a pooled dataset of all experiments as in the Sachs et al. method. We define an experiment-specific network score *S*(*G^j^* : *D_j_*) as sum (over all variables *V_i_*) of per-variable local scores *S*_local_ (*V_i_, U* : *D_j_*) of variables *V_i_*. The left part of the equation is the prior probability assigned to the choice of set U as parents of *V_i_*, and the right part is the probability of the data integrated over every possible parameterizations (*θ*) of the distribution.

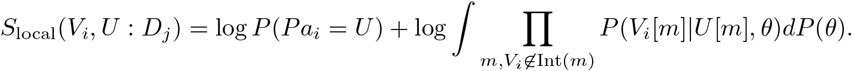

### Structure learning

Because our method uses local stochastic search (Tabu), we create an ensemble of n random starting DAGs (stored as `randomNet`, see Algorithm 1) using the procedure `createRandNet`. Empirically, we have found that *n* = 100 is adequate for the network multivariate datasets that we analyzed in this work to demonstrate empirical performance of our method (see Results). From each DAG in `randomNet`, we then search for an optimal network model using the Tabu search algorithm [10] and store the *n* networks in a list `Net`. The list `intv` of known targets is passed as an argument which incorporates interventions in the search algorithm by preventing the arcs to be incident on the targets. Next, we measure the probabilistic arc strength and direction (using the procedure `arcStrength`) for each arc as its empirical frequency given the list of networks in `Net`. We average the arc strengths for every directed arc over the networks in which corresponding target node was not intervened and store them as a list `arcProb`.

### Combining results from experiments

Given arc information (in `arcProb`, see Algorithm 1) from each experiment, we average their strengths and directions over the number of experiments where the given arc is valid (using procedure `avgNetwork`). Finally, we compute the averaged arc strengths as `avgArcs` and threshold on arc strength (using a predefined `Threshold`) in order to produce the final DAG (using procedure `learnDAG`). We found that our method performs best at a 50% threshold. We implemented “Learn and Vote” in the R programming language, making use of the `bnLearn` package [25].

#### Datasets that we used for empirical performance analysis

From six published networks, we obtained nine datasets (with associated ground-truth networks) that we analyzed in this work. To avoid bias, from each network we used both observational and interventional datasets. For synthetic networks, as observations, we drew random samples. As interventions, we set some target nodes to fixed values. Next, in order to model uncertainty, we also set one or more of the target’s children to different values (like “fat-hands”[7]) which are assumed to be unknown. Finally, we sampled data from each of the mutilated networks [26]:

- **Lizards:** a real-world dataset having three variables representing the perching behaviour of two species of lizards in the South Bimini island [27]. We generated one observational dataset and two interventional datasets from the lizards network.
- **Asia:** a synthetic network of eight variables[28] about occurrence of lung diseases and their relation with visits to Asia. For our empirical study, we created two mutilated networks: `Asia_mut1` has one observation and one interventional dataset, and `Asia_mut2` has one observational and two interventional datasets.
- **Alarm:** a synthetic network of thirty seven variables representing an alarm messaging system for patient monitoring [29]. For our study, we created two mutilated networks: `Alarm_mut1` has three observational and six interventional studies, and `Alarm_mut2` has five observational and ten interventional datasets.
- **Insurance:** a synthetic network of twenty seven variables for evaluating car insurance risks [30]. We created two mutilated networks: `Insurance_mut1`, from which we obtained one observational and five interventional datasets; and `Insurance_mut2`, from which we obtained three observational and eight interventional datasets.
- **gmInt:** a synthetic dataset containing a matrix of observational and interventional data from eight Gaussian variables, provided in the `pcalg-R` package.
- **Sachs et al.:** a cell signaling network and associated mixed observational-interventional dataset published by Sachs et al. [13], described above).

#### Causal network learning methods that we compared to “Learn and Vote”

Using the aforementioned networks and datasets, we compared the accuracy of “Learn and Vote” for network inference to the following six algorithms (implemented in `R`):

- **PC:** We used the observational datasets to evaluate DAG-equivalent structures [2], and we used Fisher’s *z*-transformation conditional independence test (varying α from 0 to 1).
- **GDS:** This is a greedy search method [15] to estimate Markov equivalence class of DAG from observational and interventional data. It works by maximizing a scoring function (*L*_0_-penalized Gaussian maximum likelihood estimator) in three phases, i.e., addition, removal and reversal of an arrow, as long as the score improves.
- **GIES:** This algorithm [15] generalizes the greedy equivalence search (GES) algorithm (Chickering 2002) to include interventional data into observational data.
- **Globally optimal Bayesian Network (simy):** This is a score-based dynamic programming approach [31] to find the optimum of any decomposable scoring criterion (like BDe, BIC, AIC). This function (simy) estimates the best Bayesian network structure given interventional and observational data but is only feasible up to about 20 variables.
- **Invariant Causal Prediction (ICP):** This method by Peters et al., [32] calculates the confidence intervals for causal effects by exploiting the invariance property of a causal (vs. non-causal) relationship under different experimental settings. We implemented it in R, making use of the `InvariantCausalPrediction` package.
- **Sachs et al. method** The Bayesian network approach used by Sachs et al. was described in Methods and Datasets above.

For each of these methods except PC, the method implementations that we used were adapted for heterogeneous datasets (see citations above).

### Performance measurement

For the purpose of quantifying the accuracies of the nine networks learned by each of the seven network algorithms, we treated the presence of an arc in the ground-truth dataset as a “positive” and its absence as a “negative”. For each inferred network and each algorithm, from the confusion matrix we computed precision, recall, and the F1 harmonic mean of precision and recall (we did not compute accuracy due to the inherent class imbalance of sparse networks), as shown in Table 1.

**Table 1.** Multi-dataset performance of “Learn & Vote” versus six other methods. Each row corresponds to a specific dataset derived from a specific underlying ground-truth network (as described in detail in Methods and Datasets). Each row is split into three performance measures (precision, recall, and the “F1” harmonic mean of precision and recall). For each sub-row, the highest performance measurement is boldfaced. Each column corresponds to a specific method for causal network inference (as described in detail in Methods and Datasets), with the performance measures of our method (“Learn and Vote”) in the rightmost column. The symbol “n/a” denotes that no performance results were available for that method on that dataset. Here, the method “simy” is only feasible for networks containing up to 20 nodes, so it failed to produce results on the larger networks. The network **size** denotes the number of nodes in the indicated network. The network **type** is as follows: RW, real-world; S, synthetic.

| Dataset | size | type | Metric | PC | GDS | GIES | ICP | simy | Sachs et al. | Learn & Vote |
| --- | --- | --- | --- | --- | --- | --- | --- | --- | --- | --- |
| Lizards | 3 | RW | Precision | **1** | **1** | **1** | 0 | **1** | **1** | **1** |
|  |  |  | Recall | **1** | **1** | **1** | 0 | **1** | 0.5 | 0.5 |
|  |  |  | F1 score | **1** | **1** | **1** | 0 | **1** | 0.667 | 0.667 |
| Asia_mut1 | 8 | S | Precision | **1** | 0.625 | 0.625 | **1** | 0.316 | 0.77 | **1** |
|  |  |  | Recall | 0.75 | 0.625 | 0.625 | 0.5 | 0.75 | **0.875** | 0.75 |
|  |  |  | F1 score | **0.857** | 0.625 | 0.625 | 0.666 | 0.444 | 0.824 | **0.857** |
| Asia_mut2 | 8 | S | Precision | **1** | 0.857 | 0.857 | **1** | 0.304 | 0.666 | **1** |
|  |  |  | Recall | 0.75 | 0.75 | 0.75 | 0.5 | **0.875** | 0.75 | 0.75 |
|  |  |  | F1 score | **0.857** | 0.8 | 0.8 | 0.666 | 0.493 | 0.706 | **0.857** |
| gmInt | 8 | S | Precision | 0.75 | 0.889 | 0.889 | **1** | 0.889 | 0.857 | **1** |
|  |  |  | Recall | 0.75 | **1** | **1** | 0.375 | **1** | 0.75 | 0.75 |
|  |  |  | F1 score | 0.75 | **0.94** | **0.94** | 0.545 | **0.94** | 0.8 | 0.857 |
| Cell signaling | 11 | RW | Precision | 0.571 | 0.419 | 0.377 | **1** | 0.422 | 0.68 | 0.89 |
|  |  |  | Recall | 0.4 | **0.9** | 0.85 | 0.45 | 0.95 | 0.85 | 0.89 |
|  |  |  | F1 score | 0.47 | 0.572 | 0.522 | 0.62 | 0.584 | 0.756 | **0.89** |
| Insurance_mut1 | 27 | S | Precision | 0.714 | 0.36 | 0.362 | 0.7 | n/a | **0.857** | 0.8 |
|  |  |  | Recall | 0.288 | 0.346 | 0.327 | 0.25 | n/a | **0.577** | 0.538 |
|  |  |  | F1 score | 0.411 | 0.352 | 0.343 | 0.368 | n/a | **0.689** | 0.643 |
| Insurance_mut2 | 27 | S | Precision | **0.714** | 0.355 | 0.366 | 0.64 | n/a | 0.676 | 0.686 |
|  |  |  | Recall | 0.288 | 0.423 | 0.423 | 0.21 | n/a | 0.442 | **0.461** |
|  |  |  | F1 score | 0.411 | 0.386 | 0.392 | 0.316 | n/a | 0.535 | **0.552** |
| Alarm_mut1 | 37 | S | Precision | 0.666 | 0.25 | 0.26 | **0.7** | n/a | 0.625 | 0.564 |
|  |  |  | Recall | 0.434 | 0.217 | 0.26 | 0.26 | n/a | **0.446** | 0.4 |
|  |  |  | F1 score | **0.526** | 0.232 | 0.26 | 0.38 | n/a | 0.52 | 0.468 |
| Alarm_mut2 | 37 | S | Precision | 0.666 | 0.411 | 0.513 | 0.6 | n/a | 0.725 | **0.769** |
|  |  |  | Recall | 0.434 | 0.456 | 0.434 | 0.21 | n/a | 0.63 | **0.642** |
|  |  |  | F1 score | 0.526 | 0.432 | 0.47 | 0.311 | n/a | 0.675 | **0.7** |

## Results

### Effect of interventions on network inference

Based on prior studies suggesting that incorporating data from interventional studies improves network inference (see Introduction), we re-analyzed the Sachs et al. [13] biological cell signaling dataset (for which a ground truth network was published [13]) using their published inference approach twice, first using observational samples only (Figure 4a) and then using an equal number of samples comprising 50% observational and 50% interventional data (Figure 4b). We found that sensitivity for detecting cell signaling interactions increases when data from observational and interventional experiments are co-analyzed (Fig. 4b), versus when only data from observational experiments are used (Fig. 4a). These results illustrate the benefit of using data from interventional experiments for causal network reconstruction.

**Fig 4.**
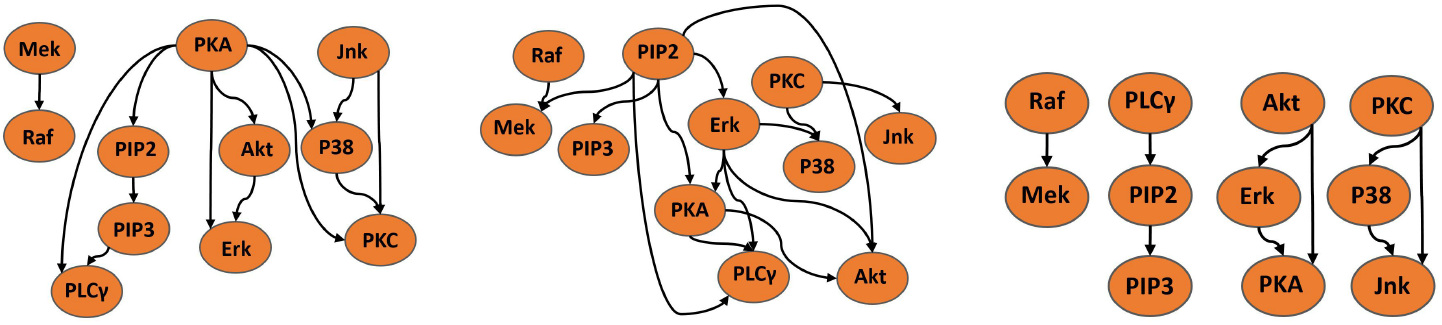
Networks inferred by (a) pooling data from two observational experiments; (b) pooling data from an observational (anti-CD3/CD28) and an interventional experiment (AKT inhibitor); and (c) “Learn and Vote” using the same experiments as in the middle panel. The structure learning statistics used are True Positive (TP), False Positive (FP) and False Negative (FN). False positives are reduced by avoiding pooling.

### Effect of pooling on network inference

Based on prior studies suggesting that pooling data from multiple experiments can lead to errors in network learning (see Introduction), we analyzed the same cell signaling dataset as in Fig. 4b, using the “Learn and Vote” method, in which data are not pooled. Compared to the the Sachs et al. inference method which was based on data pooling (Fig. 4b), use of “Learn and Vote” significantly reduced false positives, while increasing the overall robustness of the network learning (Fig. 4c).

### Systematic comparative studies

To study the performance characteristics of “Learn and Vote” for a broader class of network inference applications, we carried out a systematic, empirical comparison our method’s performance with six previously published causal network learning methods using nine datasets (from six underlying networks of small to medium size, as described above in Methods and Datasets), spanning a variety of application domains.

#### Networks learned by the seven methods on the cell signaling dataset

On the Sachs et al. dataset, the consensus networks that each algorithm learned are shown in Fig. 5a-g; the networks varied significantly in terms of density, with GDS, GIES, and simy giving large numbers of edges, and PC and ICP giving relatively sparse networks (with the PC network having many ambiguous arc directions). For each of the methods, we tabulated the numbers of correct and incorrect (or missing) arcs in the consensus networks learned (Fig. 5h). The greedy algorithms (Fig. 5b,c) and simy (Fig. 5e) are able to find most of the true positive arcs at the cost of a large number of false positives. The consensus “Learn and Vote” network (Fig. 5g) improved over the consensus network obtained using the Sachs et al. inference method (Fig. 5f), by eliminating six false positive edges and gaining a true positive edge (*PIP2* → *PKC*) (Fig. 5h, rightmost two columns). We further note that two of the putatively false interactions that were detected by “Learn and Vote”, (*P*38 → *pjnk*) and (*PKC* → *p*44.42), on further study are likely real interactions according to PCViz (www.pathwaycommons.org/pcviz) and PubMed (www.ncbi.nlm.nih.gov/pubmed). Moreover, our method had the lowest number of false positives among all seven methods and was tied for second-highest in terms of the number of true positives (Fig. 5h).

**Fig 5.**
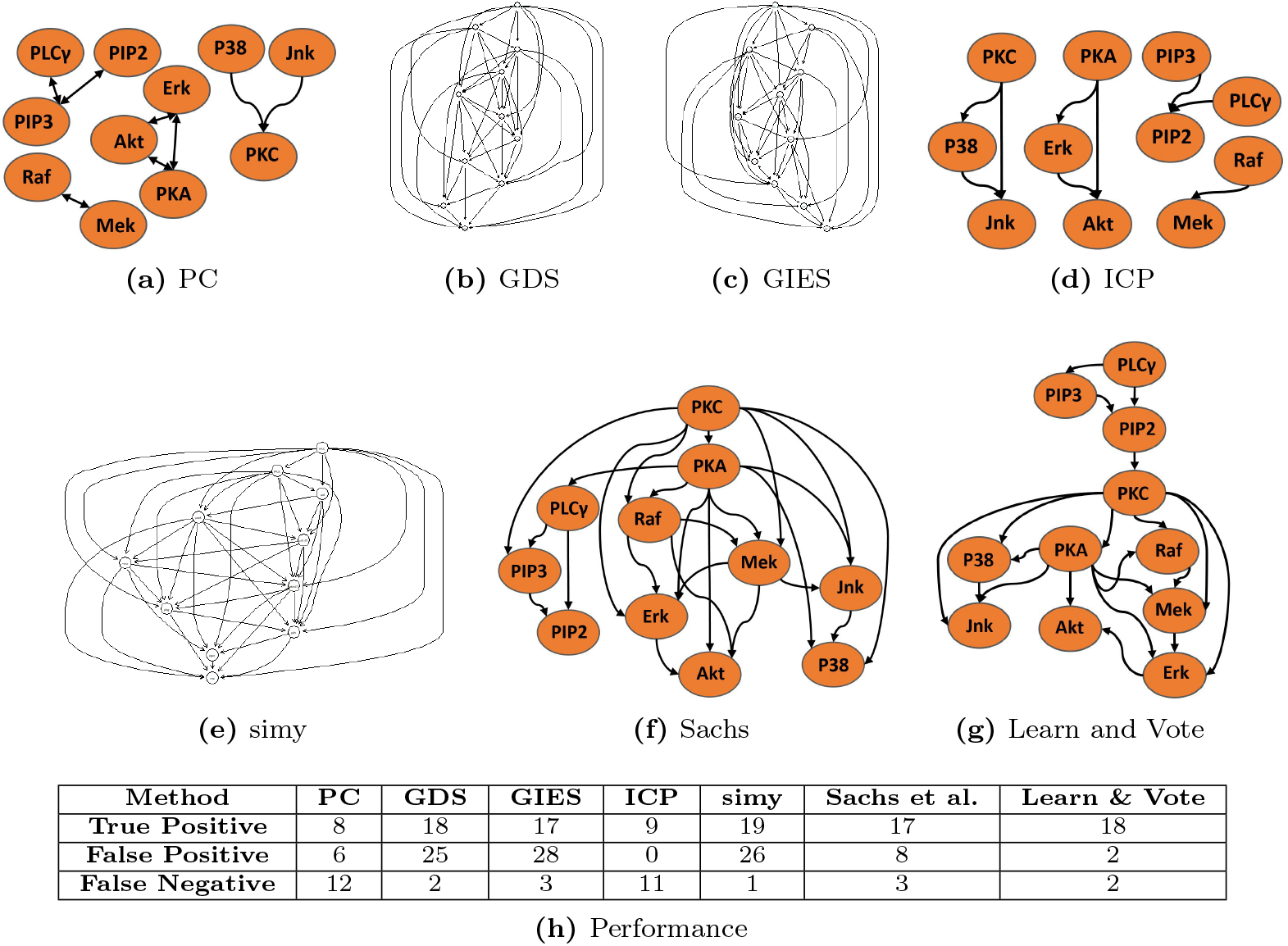
Consensus networks inferred from various algorithms (a-g) on the Sachs et al. cell signaling dataset. A bidirectional arrow between two nodes denotes that an interaction is predicted between the two nodes, but the direction of causality is ambiguous. In the table (h), each row corresponds to a component of the confusion matrix (true positives, false positives, and false negatives), and each column corresponds to a causal network inference method.

### Quantifying performance of seven network learning algorithms

In Table 1, we summarize the performance, in terms of network learning precision, recall, and F1 score of the seven network inference methods applied to nine datasets (with associated ground-truth networks) that were described in Methods and Datasets. In terms of F1 accuracy, while the PC algorithm (which used *observational* measurements) has strong performance on smaller networks, “Learn and Vote” has superior performance for learning the structure of larger networks. More broadly, “Learn and Vote” outperformed the other six algorithms in five out of nine studies in terms of precision, with the ICP method having second best performance. The positive predictive rate of our approach is higher for small or medium sized networks (i.e., fewer than 20 nodes) but decreases as the size of the network increases. In contrast, the greedy algorithms (GDS, GIES) perform well for smaller networks but suffer from lower precision on larger networks. In terms of F1, our approach outperformed the others in five out of nine studies and is more stable even when the network size increases. For very small networks (i.e., fewer than 10 nodes), the PC-based approach has good sensitivity, however, it leaves many of the arc directions ambiguous (Fig. 5a).

### Sensitivity to threshold

To study the sensitivity of our results to the threshold parameter (which was set to 0.5) for predicting a causal arc, we compared the performance of “Learn and Vote” to that of the Sachs et al. method on three different network datasets (cell signaling, Asia_mut1, and Asia_mut2; see Methods and Datasets) by plotting the sensitivity versus false positive error rate (FPR) for various threshold values (Fig. 6a). On all three datasets, in terms of area under the sensitivity-vs-FPR curve, “Learn and Vote” has a higher score than the Sachs et al. method, with the most significant performance gap occurring at thresholds where the specificity is in the range of 0.7–0.9.

**Fig 6.**
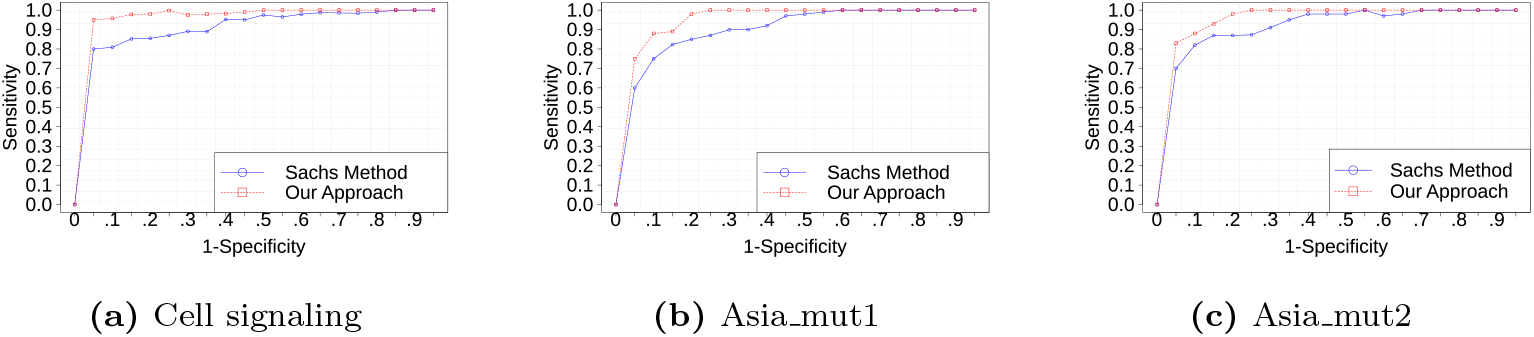
Sensitivity vs. FPR for “Learn and Vote” and the Sachs et al. method on three datasets: (a) Sachs et al. cell signaling; (b) Asia lung disease (mut1); and (c) Asia lung disease (mut2). The line plots are nonmonotonic due to the use of different random initial DAGs for different points on the line plot.

### Effect of sample size

It seems intuitive that in cases where single-experiment sample sizes are very small, separately analyzing data from individual experiments would be expected to perform poorly relative to a pooling-based approach like the Sachs et al. method. To test this, we analyzed the how the relative performances of “Learn and Vote” and the Sachs et al. method vary with sample size on the Sachs et al. dataset (for which the Sachs et al. method was specifically developed). We sampled equal numbers of data points from each experiment to prevent bias towards a particular experiment. Fig. 7 shows the performance of our method versus the Sachs et al. method by varying the numbers of samples used from each experiment. When the number of samples per experiment is very small, learning from pooled data gives a better result. For the Asia network, which has eight nodes, when the number of samples per experiment is very small (e.g., 20 samples), the performance of “Learn and Vote” is no better than the pooling-based Sachs et al. method (Fig. 7b-c). Hence, when only a small amount of data are available it is a good idea to combine them irrespective of experimental conditions. However, for large enough sample size, we see that pooling degrades accuracy of network reconstruction.

**Fig 7.**
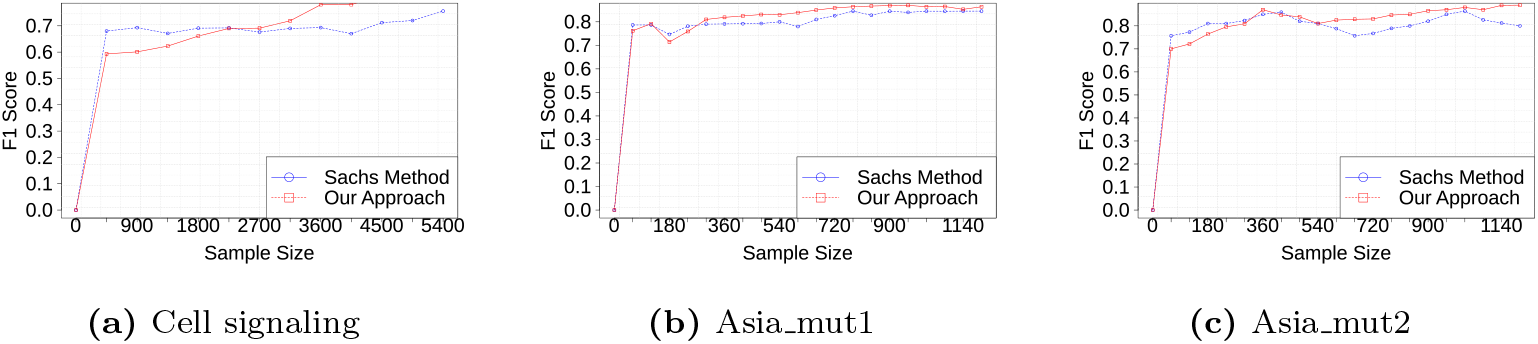
F1 vs. sample size for Learn and Vote and the Sachs et al. method, for three datasets.

## Discussion

Taken together, our results (Fig. 5 and Table 1) suggest that for analyzing datasets from studies that have imperfect interventions, greedy analysis methods (e.g., GDS, GIES) are not as accurate as “Learn and Vote”. On the other hand, ICP is conservative due to its strict invariance property and helps reduce false causal arcs to a great extent, but at the cost of sensitivity (Fig. 5d). The relatively poor performance of the PC method on the Sachs et al. dataset likely reflects the fact that it does not utilize interventional data. In future work, we plan to study the case of handling uneven samples of data from different experiments. We also plan to extend the work by choosing which interventional target is more informative in an unknown network structure. Another improvement of our approach is to see how choosing the number of random DAGs (we have taken 100) scales with network size. For example, in case of larger graphs, 100 might not be sufficient while in smaller graphs it could be overkill. One possible improvement to “Learn and Vote” would be an adaptive method for selecting the number of random initial DAGs; this is an area of planned future work.

## Conclusion

We report a new approach, “Learn and Vote,” for learning a causal network structure from multiple datasets generated from different experiments, including the case of hybrid observational-interventional datasets. Our approach assumes that each dataset is generated by an unknown causal network altered under different experimental conditions (and thus, that the datasets have different distributions). Manipulated distributions imply manipulated graphs over the variables, and therefore, combining them to learn a network might increase statistical power but only if it assumes a single network that is true for every dataset. Unfortunately, this is not always the case under uncertain interventions. Our results are consistent with the theory that simply pooling measurements from multiple experiments with uncertain interventions leads to spurious changes in correlations among variables and increases the rate of false positive arcs in the consensus network. In contrast, our “Learn and Vote” method avoids the problems of pooling by combining experiment-specific weighted graphs. We compared “Learn and Vote” with six other causal learning methods on observational and interventional datasets having uncertain interventions. We found that for most of the larger-network datasets that we analyzed, “Learn and Vote” significantly reduces the number of false positive arcs and achieves superior F1 scores. However, for cases where sample size per experiment is very small, we found that pooling works better. Our findings (i) motivate the need to focus on the uncertain and unknown effects of interventions in order improve causal network learning precision, and (ii) suggest caution in using causal learning algorithms that assume perfect interventions, in the context of real world domains that have uncertain intervention effects.

## Acknowledgment

Research reported in this publication was supported in part by the National Center for Advancing Translational Sciences (NCATS), National Institutes of Health, through the Biomedical Data Translator program (award OT2TR002520 to SAR). The content is solely the responsibility of the authors and does not necessarily represent the official views of the National Institutes of Health.

